# G-protein coupled receptor activity mediates detrusor smooth muscle phasic contractility through regulation of membrane potential

**DOI:** 10.64898/2026.08.10.743960

**Authors:** Jason L. Rengo, Thomas J. Heppner, Grant W. Hennig, Nicholas R. Klug, Sloane Stamp, Mark T. Nelson, Gerald M. Herrera

**Author notes:** Corresponding Author: Gerald M. Herrera, Ph.D., Department of Pharmacology Larner College of Medicine University of Vermont, 149 Beaumont Ave. FMRB 461, Burlington, VT 05405.

## Abstract

The urinary bladder functions to store and release urine, yet how the sensation of bladder fullness is conveyed and perceived to the central nervous system is not understood. During bladder filling, the detrusor smooth muscle (DSM) generates phasic contractions, resulting in pressure fluctuations within the bladder. These transient pressure events drive bursts of afferent nerve activity, yet the underlying mechanism leading to rhythmic contractions remains unclear. Here, we examined the role of Gq protein-coupled receptor (GqPCR) activity on DSM excitability and contractility. Using ex vivo pressurized urinary bladder preparations and sharp microelectrode experiments on bladder strips from mice, we evaluated whole bladder transient pressure events, whole bladder DSM Ca^2+^ activity, and membrane potential in bladder strips. We found that global inhibition of urinary bladder GqPCR activity with YM-254890 abates phasic contractility and transient pressure events through a reduction in DSM Ca^2+^ activity and propagation of Ca^2+^ waves. Further, we found inhibition of GqPCR significantly hyperpolarizes DSM, reducing action potentials and decreasing excitability, and activation of protein kinase C restores membrane potential to baseline levels. These findings highlight that GqPCR activity mediates DSM excitability and contractility in such a way as to result in phasic detrusor contractions and transient pressure events.

## INTRODUCTION

The main function of the urinary bladder is to store urine until the need to void arises. While seemingly a simple process, this requires coordination of sensory, cognitive, and motor systems such that voiding is neither overly frequent, nor is urine retained too long (Andersson & Arner, 2004). The bladder wall is comprised of detrusor smooth muscle (DSM) cells and it is the contraction and relaxation of DSM that mediates the voiding and storage functions of the bladder. During filling, the sympathetic nervous system relaxes the bladder via beta adrenergic receptors (Andersson & Arner, 2004). During voiding, the parasympathetic nervous system initiates contraction of the DSM predominantly via M3 muscarinic Gq protein-coupled receptors (GqPCR) while the somatic nervous system allows voluntary control of the external urethral sphincter (Andersson & Arner, 2004). While the pathways controlling the storage and voiding phases of micturition are well known, the underlying physiological mechanism of how bladder fullness is sensed is less clear. DSM is an excitable tissue capable of exhibiting action potentials (Heppner *et al*., 1997; Hashitani & Brading, 2003a, b). These action potentials can be coordinated and propagated in waves throughout the bladder to produce local phasic contractions that result in pressure fluctuations, termed transient pressure events (TPE). The rate of rise in these pressure transients correlates with the amplitude of coincident bursts of afferent nerve activity to the central nervous system, suggesting a causative role for TPE in signaling bladder fullness (Heppner *et al*., 2016). However, it is unclear what drives phasic contractions. The rhythmic nature of TPE suggests some form of intrinsic activation and coordination across DSM. While voiding contractions result from the pelvic nerve releasing the neurotransmitters adenosine triphosphate (ATP) and acetylcholine (ACh) onto the bladder, purinergic and muscarinic GqPCR appear unrelated to the generation of phasic contractions and TPE during bladder filling (Herrera *et al*., 2000; Imai *et al*., 2001). Further, TPE are abolished by blocking L-type voltage-dependent calcium channels (VDCCs) but are unaffected by inhibiting neural activity with the voltage-dependent Na^+^ channel blocker, tetrodotoxin, suggesting that phasic contractility is intrinsic to the DSM (Herrera *et al*., 2000; Imai *et al*., 2001). Activation of GqPCR increases intracellular calcium (Ca^2+^) and protein kinase C (PKC) activity, both of which can regulate potassium (K^+^) and Ca^2+^ channels, leading to changes in DSM excitation-contraction coupling (Herrera *et al*., 2000, 2001). Thus, GqPCR are prime candidates to influence phasic DSM contractility and TPE. GqPCR expression is ubiquitous within the bladder, including prostaglandin, thromboxane, serotonin, bradykinin, neurokinin, endothelin, histamine, and numerous other receptors, suggesting one or more of these receptor classes may be involved in DSM phasic contractility (Regard *et al*., 2008).

The GqPCR signaling cascade results in the activation of phospholipase C, which in turn cleaves membrane bound phospholipid phosphatidylinositol 4,5-bisphosphate into 1,4,5-trisphosphate (IP3) and diacylglycerol (DAG). Downstream effects include IP3 mediated Ca^2+^ release from intracellular stores and activation of protein kinase C (PKC) via DAG and Ca^2+^ (Jiang *et al*., 2022). PKC activity regulates K^+^ channel open probability in many types of smooth muscle tissue, including DSM. PKC decreases large-conductance calcium-activated potassium channel (BK) open probability via phosphorylation of serine residues S695 and S1151 on the C-terminus (Zhou *et al*., 2010). PKC activity can also modulate bladder smooth muscle phasic contractility and inhibit BK currents (Hristov *et al*., 2014). PKC may also reduce voltage-gated K^+^ channel (Kv) and small-conductance calcium-activated K^+^ channel (SK) current density (Jackson *et al*., 2016; Xing *et al*., 2024). Collectively, PKC activity could be expected to inhibit K^+^ channels, leading to membrane depolarization, Ca^2+^ entry through VDCC, and DSM phasic contractility.

Here, we hypothesize GqPCR activity regulates phasic contractions and resultant TPE through modulation of DSM membrane potential (V_M_). Using ex vivo pressurized urinary bladder preparations and sharp microelectrode experiments on DSM strips from mice, we examined the effect of GqPCR inhibition on DSM contractility and excitability. We used the small molecule GqPCR inhibitor YM-254890 to probe the function of GqPCR in mediating TPE. YM-254890 blocks GqPCR activation by inhibiting GTP to GDP exchange on the intracellular domain (Nishimura *et al*., 2010). The inhibitory effects are due to Gq isoforms adopting conformations similar to the YM-254890 bound state, which stabilize an inactive GDP-bound form (Trent *et al*., 2025). We found that global inhibition of urinary bladder GqPCR abolishes phasic contractility and TPE through a reduction in DSM Ca^2+^ activity and its spread across the bladder wall. Further, we investigated the link between GqPCR activity and DSM V_M_. We found the effects of GqPCR activity on contractility require an electrochemical K^+^ driving force and inhibition of GqPCR activity significantly hyperpolarizes DSM, reducing action potentials and decreasing excitability. Taken together, these findings suggest that GqPCR activity mediates DSM excitability and contractility in such a way as to result in phasic detrusor contractions and TPE.

## RESULTS

### Spontaneous Bladder Contractility Requires GqPCR Activity

To examine the contribution of GqPCR activity to contractility in whole urinary bladders, ex vivo bladders were cannulated and attached to a syringe pump and inline pressure transducer. Urinary bladders from C57bl/6J mice were partially filled and TPE dynamics were recorded over 5-min intervals (Heppner *et al*., 2024) (Figure 1A & B).

**Figure 1.**
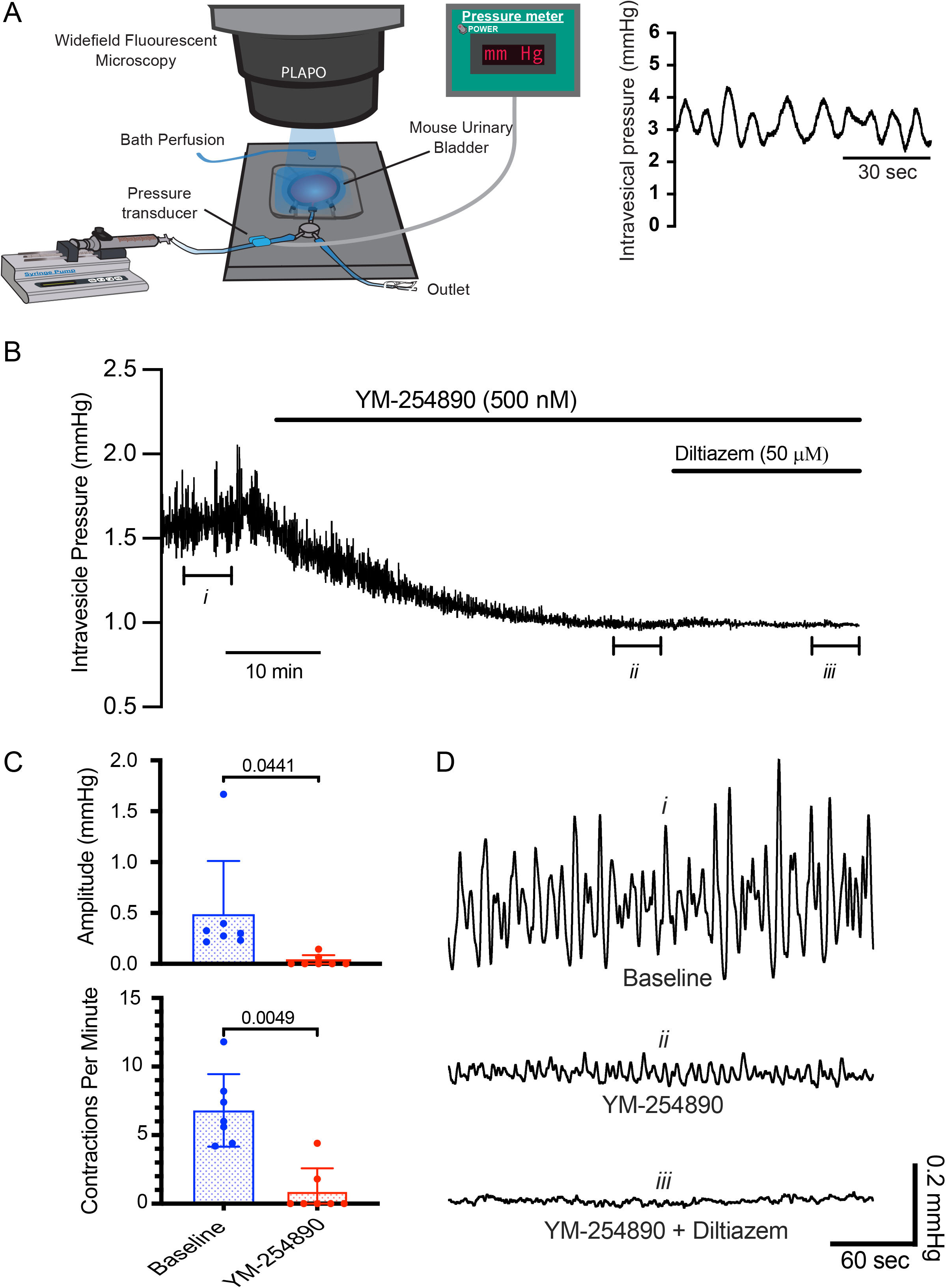
Inhibition of GqPCR activity reduces whole bladder transient pressure events. A, diagram of the ex vivo pressurized bladder set-up (adapted from Herrera et al 2026). The bladder is partially filled with physiological saline solution (PSS) to induce transient pressure events and phasic contractility is monitored with an inline pressure transducer. B, representative trace of a pressurized whole bladder preparation showing the effects of the GqPCR inhibitor YM-254890 (500 nM) and L-type voltage dependent Ca^2+^ channel blocker diltiazem (50 μM). C, summary data of the effect of YM-254890 on transient pressure events compared to PSS (baseline condition). D, representative pressure traces from each condition shown in (B). Data are mean ± SD; statistical analysis was performed with a paired t-test from N = 7 bladders.

After filling urinary bladders with physiological saline solution (PSS), intravesical pressure and TPE were allowed to stabilize before making any pharmacological interventions. The ensuring period in which intravesical pressure and TPE properties were analyzed before exposing the preparations to pharmacological treatments was defined as the “baseline” condition in all experiments. At baseline, TPE amplitude was 0.49 ± 0.52 mmHg and frequency was 6.8 ± 2.6 contractions per minute. Inhibition of GqPCR signaling with YM-254890 (500 nM), decreased mean TPE frequency (87%, p=0.0049) and amplitude (94%, p=0.0441) (n=7 bladders, Figure 1C), supporting the hypothesis that GqPCR activity is necessary to produce DSM phasic contractions. Activation of L-type VDCC has previously been shown to regulate DSM excitability. Here, subsequent application of the VDCC blocker diltiazem (50 μM) inhibited any residual phasic activity (0.0 ± 0.0 contractions per minute) (Figure 1D). These results suggest that GqPCR activity provides a major stimulatory signal driving phasic bladder contractions.

### GqPCR-Dependent Bladder Contractions are Independent of Neurotransmitter Release

The endogenous neurotransmitter ACh is a potent contractile agent in DSM, acting via the GqPCR M3 muscarinic receptors. To rule out the possibility that the endogenous neurotransmitter ACh released from nerve endings within the bladder wall is generating TPE, we applied the muscarinic antagonist atropine (50 μM). Muscarinic receptor inhibition had no effect on TPE amplitude (0%, p=0.8053) and frequency (-7%, p=0.1069) compared to baseline (n = 4 bladders, Figure 2). To confirm efficacy of atropine, we then added the muscarinic agonist carbachol (CCh, 10 μM) in the presence of atropine to the perfusion solution, which had no effect on TPE amplitude (0%, p=0.8795) or frequency (-2%, p=0.6329) compared to baseline (Figure 2). Furthermore, we and others have previously demonstrated the neuroinhibitory compound tetrodotoxin does not alter TPE but does inhibit EFS induced contractions (Herrera *et al*., 2000; Imai *et al*., 2001). Together, these data indicate generation of phasic contractions and TPE are not mediated by transmitters released by nerves. However, this finding does not exclude a role for agonists acting through GqPCR released from other cell types.

**Figure 2.**
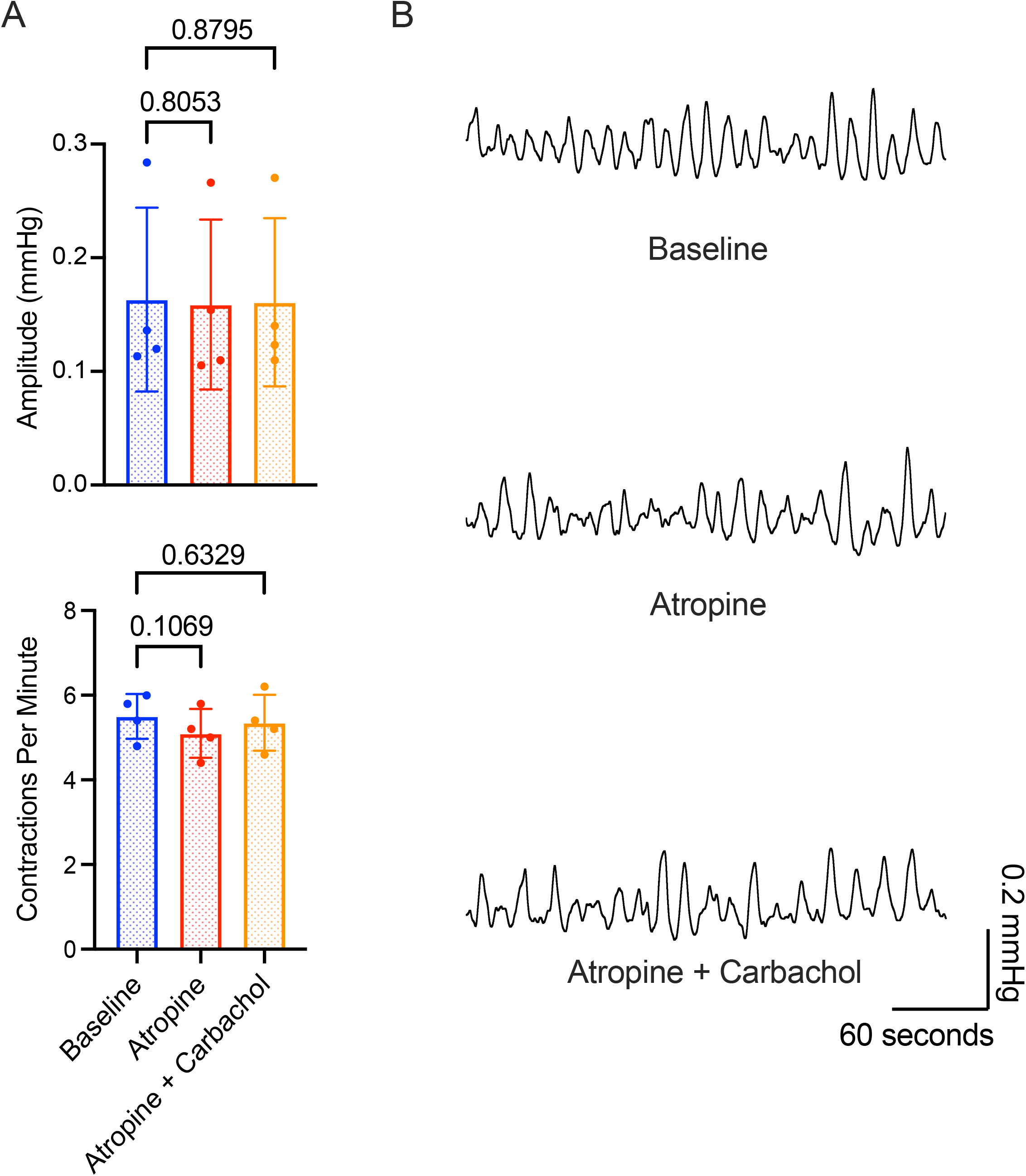
The muscarinic receptor antagonist atropine has no effect on urinary bladder transient pressure events (TPE). A, summary data showing inhibition of muscarinic receptors with atropine (50 μM) did not alter TPE compared to physiological saline solution (baseline condition). Subsequent addition of the muscarinic receptor agonist carbachol (10 μM) had no effect on TPE, confirming the efficacy of atropine. B, representative pressure traces from each condition shown in (A). Data are mean ± SD and statistical analysis was performed with repeated measures ANOVA with Dunnett’s test for multiple comparisons from N = 4 bladders.

### Pharmacological Analysis Demonstrates GqPCR-Specific Actions of YM-254890

The muscarinic agonist CCh enhances DSM contractility in the ex vivo pressurized bladder preparation through activation of M3 GqPCR (Andersson & Arner, 2004). To test the efficacy of YM-254890 on a known GqPCR response, we applied CCh (10 μM) in the presence and absence of YM-254890. In the absence of YM-254890, CCh induced a contraction of 17.13 ± 10.58 mmHg. YM-254890 pre-treatment resulted in a 92% decrease in the CCh-induced contraction to 1.30 ± 1.04 mmHg (p = 0.0143, n = 6 bladders, Figure 3A & B). These results demonstrate that YM-254890 inhibits a GqPCR signaling response evoked via M3 muscarinic receptor stimulation in the ex vivo bladder preparation.

**Figure 3.**
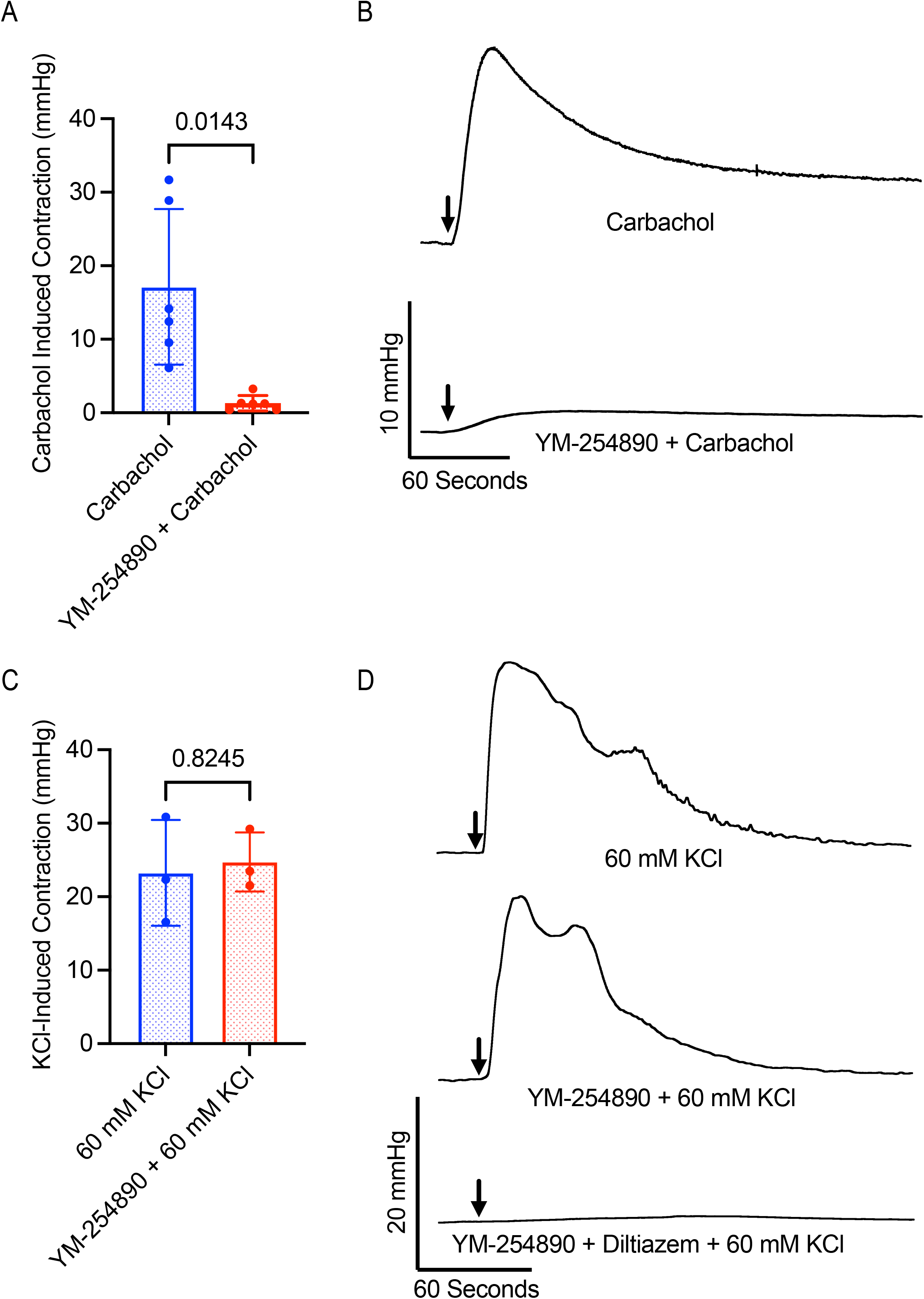
YM-254890 blocks the effect of the known bladder smooth muscle GqPCR agonist carbachol but not voltage-dependent calcium channel activation by high concentration potassium (KCl). A, summary data showing the GqPCR inhibitor YM-254890 (500 nM) blocks whole bladder contractions induced by the GqPCR agonist carbachol (10 μM) in N = 6 bladders. B, representative pressure traces from each condition shown in (A). Arrows indicate the addition of carbachol. C, summary data showing KCl (60 mM) induced contractions in the presence and absence of YM-254890 (500 nM) from N = 3 bladders. D, representative pressure traces from summarized data shown in (C). The L-type voltage dependent Ca^2+^ channel antagonist diltiazem (50 μM), but not YM-254890, blocks KCl induced contractions in whole bladder preparations. Arrows indicate the addition of 60 mM KCl. Data are mean ± SD and statistical analysis was performed with a Welch’s t-test (for A) or paired t-test (for C).

Given the fundamental importance of VDCC in regulating DSM contractility (Figure 1) (Heppner *et al*., 1997; Hashitani *et al*., 2004), another possible explanation for YM- 254890 inhibition of TPE is that YM-254890 is inhibiting the activity of VDCC. To test for this possibility, we applied high concentration potassium chloride (60 mM KCl), which depolarizes the cell membrane of DSM, activating VDCC and eliciting smooth muscle contraction (Nelson & Worley, 1989). In the absence of YM-254890, introduction of 60 mM KCl into the bath solution resulted in a contraction of 23.25 ± 7.19 mmHg. In the presence of YM-254890, KCl induced a contraction of 24.74 ± 4.01 mmHg (p=0.8245) (n = 3 bladders, Figure 3C & D). This result is consistent with previous studies showing YM-254890 has no effect on agonist-independent contractions induced by high KCl (Tamalunas *et al*., 2022). Application of KCl in the presence of diltiazem (and continued presence of YM-254890) did not elicit a contractile response, demonstrating that KCl-induced contraction is attributed to the activity of VDCC (0.19 ± 0.22 mmHg, Figure 3D). These results suggest that YM-254890 inhibits TPE independent of VDCC activity.

### GqPCR Signaling Promotes Coordinated DSM Ca^2+^ Activity and Wave Propagation

[Ca^2+^]_i_ is a vital component of excitation-contraction coupling in DSM, linking electrical activity (action potentials) with the contractile process via Ca^2+^ entry through VDCCs (Hashitani *et al*., 2001). Thus, monitoring DSM [Ca^2+^]_i_ with fluorescent Ca^2+^ indicators can be used as a proxy for DSM excitability (Heppner *et al*., 2005). We used mice expressing the genetically encoded Ca^2+^ indicator, GCaMP6f, in smooth muscle to monitor DSM Ca^2+^ signal patterns during bladder filling. Under normal filling conditions, Ca^2+^ signals occur sporadically throughout the bladder wall, with some areas generating large bursts of activity that spread to adjacent cells that are visualized as Ca^2+^ waves (Figure 4, Movie 1). These Ca^2+^ waves generate phasic contractions (Hashitani *et al*., 2001) and likely underlie transient pressure events seen in the ex vivo pressurized bladder and cystometry. We further assessed the spatiotemporal properties of DSM Ca^2+^ signals during bladder filling using an unbiased standard deviation method to extract DSM Ca^2+^ events to quantify the overall duration of Ca^2+^ activity sampled at every point on the bladder surface (Prevalence) and the local coordination of Ca^2+^ activity between DSM bundles to determine how this activity is synchronized across the bladder wall (Coincidence). This method was previously described by our group (Herrera *et al*., 2026). Prevalence was quantified as seconds of Ca^2+^ activity per minute occurring on the visible bladder hemisphere (%Bladder surface x s.min^-1^) (Figure 4A). Coincidence was quantified as the percent neighborhood search area (radius 1mm) containing synchronous activity and expressed as both maximum and average (Figure 4A). Each parameter was plotted as a spatiotemporal map (Figure 4A) and also presented as single values representing the overall amount (Prevalence) and average synchrony (Coincidence) of Ca^2+^ activity (Figure 4B&C). Following inhibition of GqPCR with YM-254890, Ca^2+^ prevalence decreased 92% (p=0.0337) (n = 5 bladders, Figure 4B, Movie 2). Coincidence analysis showed average synchrony amongst DSM bundles was reduced by 81% (p = 0.0216), indicating less coordination between adjacent DSM cells and a decreased spread of Ca^2+^ activity (Figure 4C).

**Figure 4.**
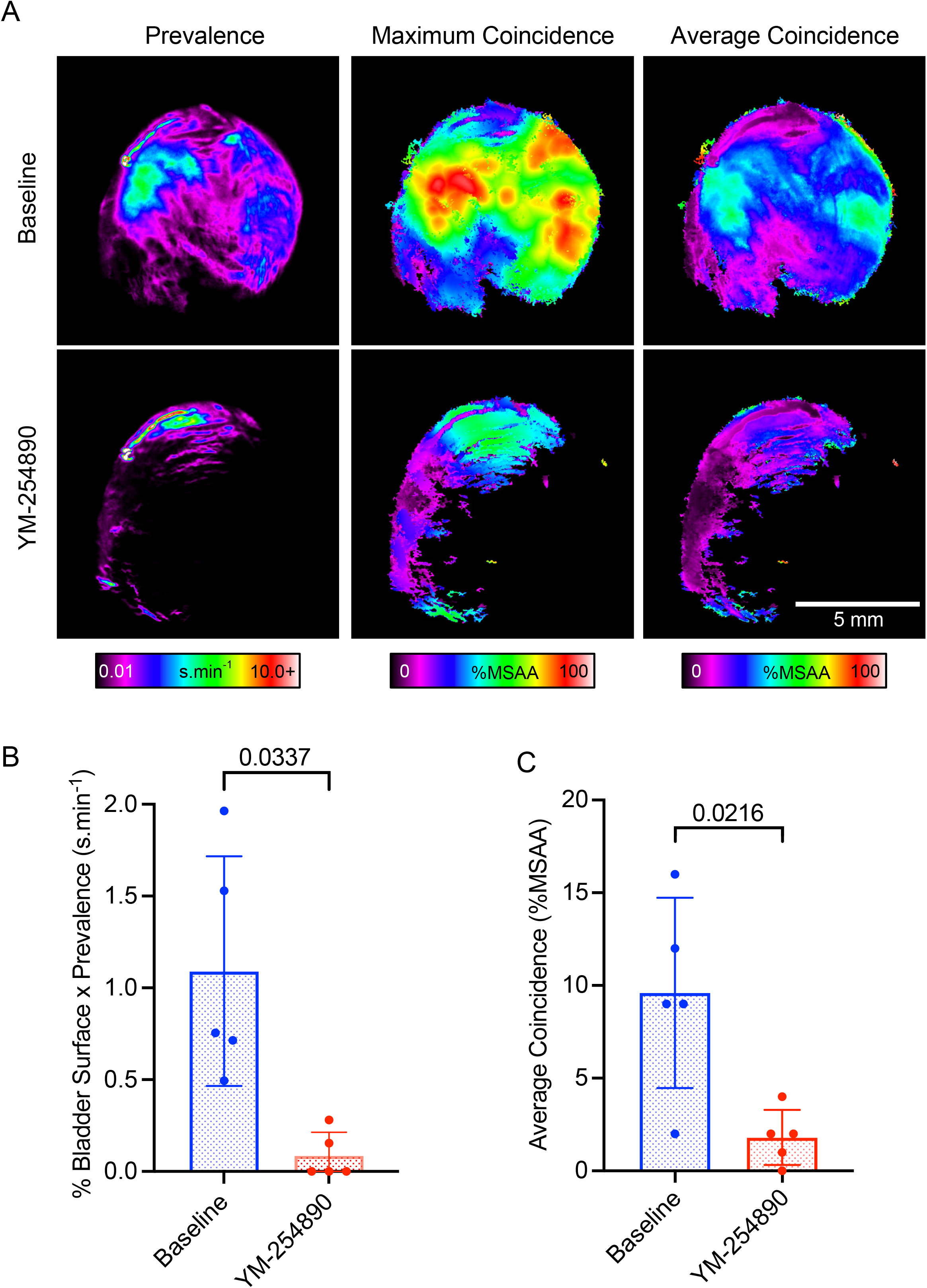
**Inhibition of GqPCR with YM-254890 reduces urinary bladder smooth muscle Ca^2+^ activity prevalence and coincidence**. A, representative experiment showing extracted Ca^2+^ events as a spatiotemporal map to quantify prevalence and coincidence of Ca^2+^ activity in urinary bladder smooth muscle. Recordings were taken using widefield microscopy over 5 minutes from mouse bladders with a fluorescent Ca^2+^ indicator specific to smooth muscle. Prevalence is presented as the total duration of Ca^2+^ activity each minute from every point of the bladder surface (%Bladder x seconds of Ca^2+^ activity per minute). Coincidence is presented as % of maximum search area activity within a radius of 1mm exhibiting simultaneous Ca^2+^ events (%MSAA). Coincidence is shown as both the peak value at every point on the bladder surface (Max%MSAA) and average coincidence (Avg%MSAA). B, summary data of Ca^2+^ activity showing prevalence was significantly reduced following application of YM-254890 (500 nM, N = 5 bladders). C, summary data of Ca^2+^ activity showing synchrony of detrusor smooth muscle cells was greatly reduced in the presence of YM-254890. Data are mean ± SD and statistical analysis was performed with a paired t-test.

### The Relaxing Effect of YM-254890 on DSM Requires an Electrochemical Driving Force for K^+^

V_M_ is a key regulator of DSM excitability and contractility, and K^+^ channels are integral to normal control of DSM V_M_ (Heppner *et al*., 1997; Hashitani & Brading, 2003a, b; Thorneloe & Nelson, 2003). Resting V_M_ of DSM cells is typically around – 40 mV. Under physiological conditions, the K^+^ equilibrium potential (E_K_) would be expected to be around – 84 mV (Hille, 1984). Thus, there is a substantial driving force for outward K^+^ movement, and slight changes in K^+^ channel activity can have a dramatic impact on DSM V_M_ (Thorneloe & Nelson, 2003). To determine whether GqPCR activity exerts effects on DSM contractility via modulation of K^+^ channel activity, we performed a physiological voltage clamp experiment where we bathed ex vivo bladders in 120mM KCl PSS, which is close to the expected intracellular concentration of K^+^ (140 mM) in smooth muscle. Thus, E_K_ and the V_M_ (0 mV) should be similar and modulating K^+^ channel activity should not affect DSM V_M_. As such, if YM-254890 alters DSM contractility by altering K^+^ channel activity, we would expect it to be without effect under these conditions, as there is no longer a driving force for K^+^ movement across the DSM cell membrane. As expected, replacing PSS (Figure 5A&C) with 120 mM KCl PSS produced a robust contraction with intravesical pressure stabilizing above baseline levels (Figure 5B). Interestingly, we found that in the presence of 120 mM KCl PSS, intravesical pressure following application of YM-254890 was unchanged (p=0.753) while subsequent inhibition of VDCC yielded a potent relaxation (n = 5 bladders, p=0.0054) (Figure 5D&E). This not only strongly supports our conclusion that YM- 254890 does not block VDCC (Figure 3C&D) but also indicates that GqPCR activity requires an electrochemical driving force for K^+^ in order to regulate contractility. To directly explore the possibility that YM-254890 alters bladder contractility by affecting DSM excitability, we next obtained direct measurements of DSM V_M_.

**Figure 5.**
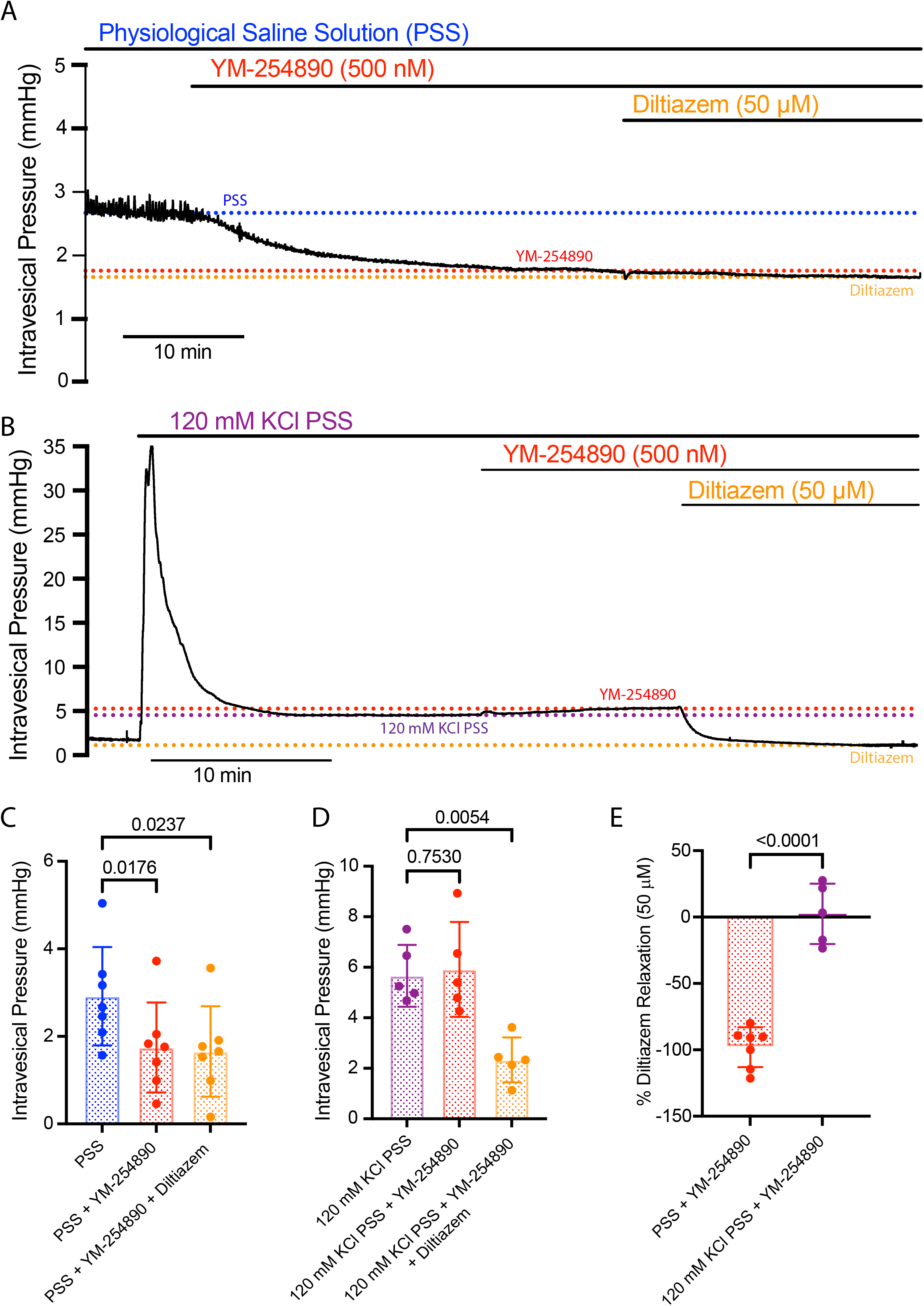
The relaxing effect of YM-254890 on bladder smooth muscle requires an electrochemical potassium driving force. A, representative trace showing the effect of YM-254890 (500 nM) on intravesical pressure. Dashed lines correspond to baseline intravesical pressure for physiological saline solution (PSS, blue), YM-254890 (red), and diltiazem (orange). B, representative trace showing the effect of YM-254890 in the presence of 120 mM KCl PSS. Superfusion of 120 mM KCl PSS eliminates the K^+^ electrochemical gradient and depolarizes bladder smooth muscle membrane potential (V_M_) close to 0 mV, increasing the open probability of L-type voltage dependent calcium channels (VDCC). Dashed lines correspond to baseline intravesical pressure for 120 mM KCl PSS (purple), YM-254890 (red), and diltiazem (orange). C, summary data showing YM-254890 relaxes urinary bladder smooth muscle to a similar extent as the VDCC blocker diltiazem (50 μM) from N = 7 bladders. D, summary data showing little effect of YM-254890 on intravesical pressure in the presence of 120 mM KCl PSS from N = 5 bladders. Subsequent application of the VDCC blocker diltiazem produces significant relaxation. E, summary data comparing the effect of YM-254890 on intravesical pressure in the presence and absence of 120 mM KCl PSS. Results are expressed as a percentage of the final intravesical pressure from the diltiazem condition. Under physiological conditions, YM-254890 produces significant urinary bladder relaxation but has little effect when V_M_ is depolarized with 120 mM KCl PSS. Data are mean ± SD and statistical analysis was performed with repeated measures ANOVA with Dunnett’s test for multiple comparisons (for C & D) or Welch’s t-test (for E).

### GqPCR Signaling Regulates DSM Contractility Through Changes in Excitability/Membrane Potential

To test the hypothesis that GqPCR activity regulates phasic contractility through changes in cell excitability, we performed sharp microelectrode electrophysiology to obtain recordings of DSM V_M_ (Figure 6ABCD). DSM exhibited characteristic action potentials. Following successful impalement of a single DSM cell, V_M_ was recorded at – 40 ± 11 mV (n = 7), consistent with previous recordings of resting V_M_ (Figure 6 D) (Heppner *et al*., 1997; Hashitani & Brading, 2003a, b; Heppner *et al*., 2005). Indeed, inhibition of GqPCR activity with YM-254890 hyperpolarized DSM by 6 mV to – 46 ± 10 mV (p=0.0352, n = 7 bladder sheets) and as expected reduced action potential frequency (Figure 6 E&F).

**Figure 6.**
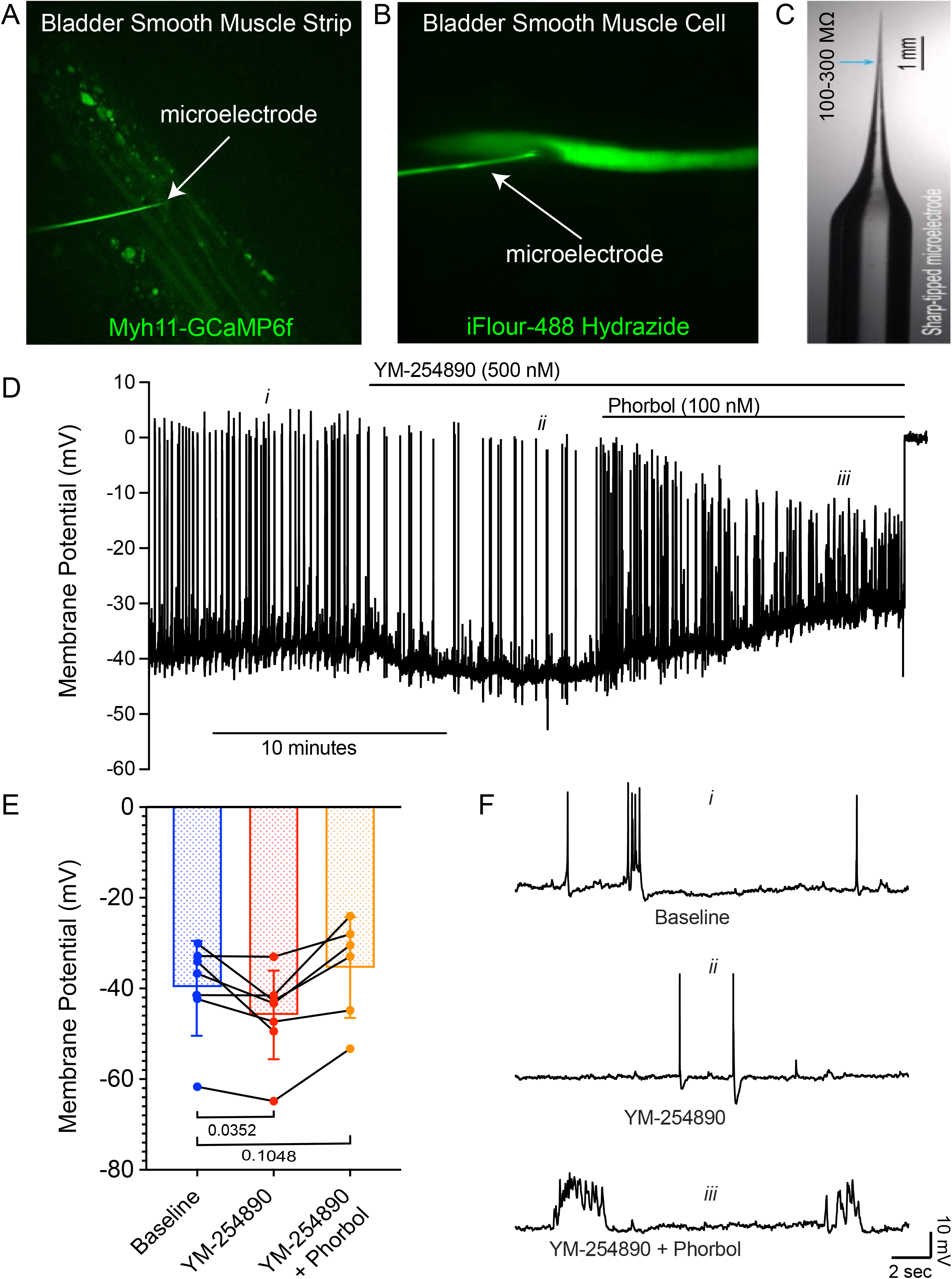
Inhibition of GqPCR with YM-254890 results in bladder smooth muscle membrane potential hyperpolarization. A, example of the sharp microelectrode experimental setup. A bladder strip with a genetic fluorescent indicator specific to smooth muscle (Myh11-GCaMP6f) was pinned in a custom chamber and a microelectrode filled with iFlour-488 Hydrazide was positioned just above the tissue. B, image showing successful impalement of an individual bladder smooth muscle cell. Following impalement, iFlour-488 Hydrazide rapidly fills the cell allowing visualization of the typical spindle-shaped morphology of bladder smooth muscle. C, example of a sharp microelectrode used to impale bladder smooth muscle. D, representative trace showing the effects of YM-254890 (500 nM) and the protein kinase C activator phorbol ester (100 nM) on bladder smooth muscle membrane potential (V_M_). E, summary data showing YM-254890 (N = 7) produces smooth muscle membrane hyperpolarization compared to baseline while subsequent addition of phorbol ester (N = 6) results in membrane depolarization. F, representative traces of V_M_ under each condition shown in (D). Data are mean ± SD and statistical analysis was performed by mixed-effects analysis with two-stage linear step-up procedure of Benjamini, Krieger and Yekutieli.

Since PKC activity lies downstream of GqPCR activation, we sought to test whether pharmacological stimulation of PKC could restore DSM V_M_ following YM-254890- induced hyperpolarization (Figure 6). We used the PKC activator phorbol 12,13- dibutyrate (phorbol ester, 100 nM). Phorbol ester caused membrane depolarization close to the original V_M_ and increased action potential frequency (-36 ± 11 mV, p=0.1048, n = 6 bladder sheets).

These results suggest that GqPCR signaling promotes excitability through PKC-dependent depolarization.

### PKC and BK Channel Modulation Partially Rescue Bladder Contractility

Our electrophysiological studies support the idea that GqPCR activity exerts an excitatory influence (depolarizing) on DSM, resulting in TPE activity. Blocking GqPCR activity with YM-254890 hyperpolarizes DSM, and this hyperpolarization could be reversed by pharmacologically stimulating PKC with phorbol ester. Thus, we sought to test the ability of phorbol ester to restore TPE contractility in the pressurized bladder in the presence of YM-254890. As already observed (Figure 1), YM-254890 greatly attenuates TPE amplitude and frequency (Figure 7 A&B). Pharmacological stimulation of PKC with phorbol ester (100 nM) results in a modest return in TPE amplitude (p=0.1324) but not frequency (p=0.0107) compared to baseline (Figure 7 A&B) (n = 7 bladders). Subsequent addition of the VDCC blocker diltiazem completely inhibited TPE (Figure 7 A&B).

**Figure 7.**
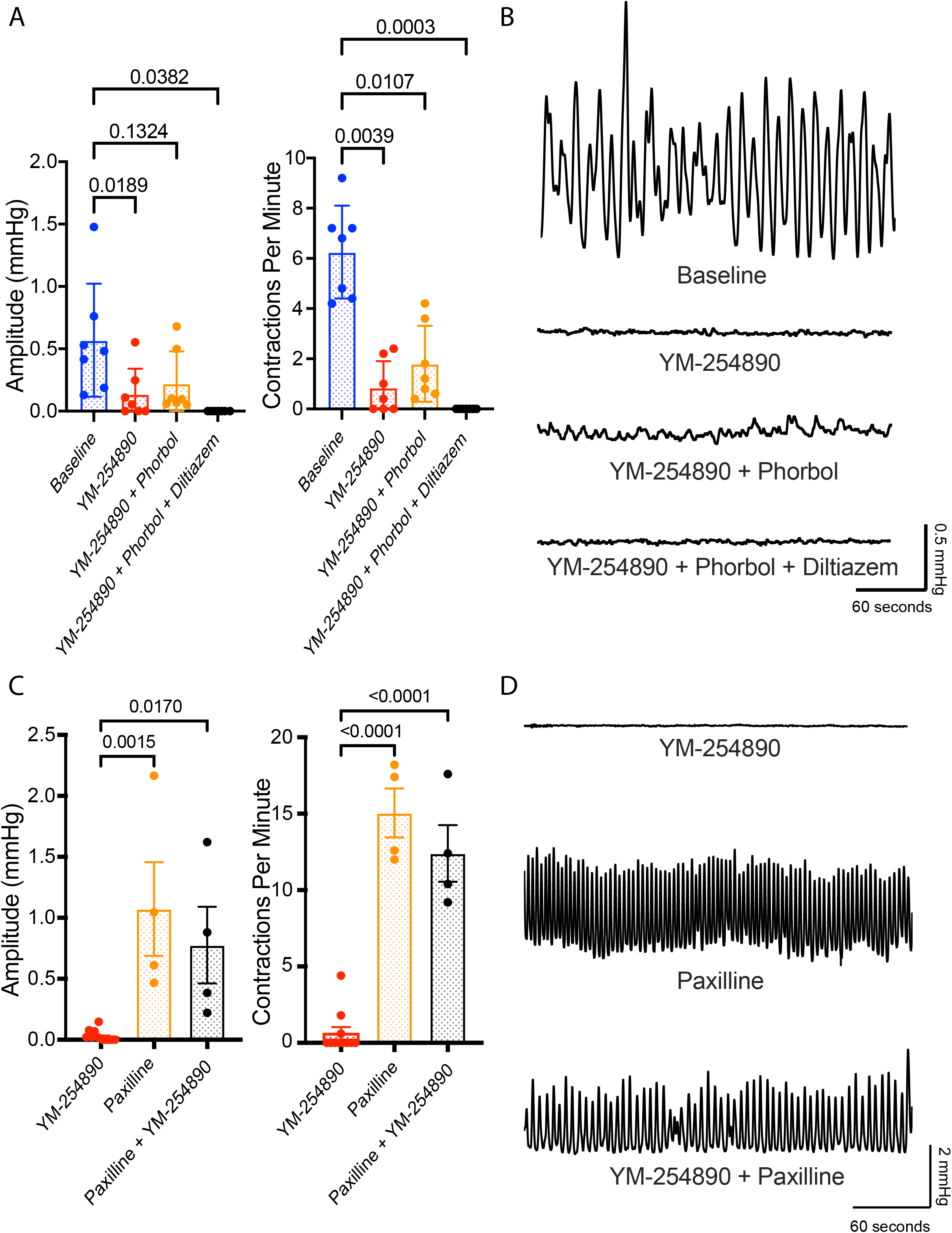
Protein kinase C and BK channel modulation partially rescue bladder contractility following GqPCR inhibition. A, summary data showing the protein kinase C activator phorbol ester (100 nM) in the presence of YM-254890 (500 nM) results in a modest return of transient pressure event amplitude, but not frequency. The addition of the L-type voltage dependent Ca^2+^ channel blocker diltiazem (50 μM) blocks phasic contractility and transient pressure events, suggesting YM-254890 and phorbol ester are acting predominantly through membrane potential rather than the smooth muscle contractile apparatus (N = 7 bladders). B, representative pressure traces from each condition shown in (A). C, the BK channel blocker paxilline (5 μM) produces significant increases in TPE amplitude and frequency. Subsequent GqPCR inhibition with YM-254890 in the continued presence of paxilline results in modest reductions that remain significantly greater when compared to YM-254890 alone (N = 4-11 bladders). D, representative pressure traces from each condition shown in (C). Data are mean ± SD and statistical analysis was performed by repeated measures (for A) or one-way ANOVA (for C) with Dunnett’s test for multiple comparisons.

This result supports our electrophysiological finding that PKC activation can restore DSM excitability after GqPCR signaling inhibition with YM-254980. Since we found that YM-254980 induced a significant DSM membrane hyperpolarization (Figure 6), a likely explanation is that GqPCR signaling may exert a PKC-dependent inhibitory influence on K^+^ conductance in DSM. One of the most important regulators of DSM excitability, which is also susceptible to inhibition by PKC, is the large-conductance calcium-activated potassium channel (BK channel) (Heppner *et al*., 1997; Schubert & Nelson, 2001; Meredith *et al*., 2004). To test the hypothesis that GqPCR signaling enhances DSM excitability through PKC-dependent inhibition of BK channels, we applied YM- 254890 in the presence of the BK channel blocker paxilline (5 μM). Inhibition of BK channels produced considerable increases in TPE amplitude and frequency, consistent with previous reports (Hristov *et al*., 2014). Subsequent GqPCR inhibition produced a modest reduction in TPE amplitude and frequency that remained significantly greater than when compared to YM-25890 alone (p=0.0170 and p<0.0001 respectively, n = 4- 11 bladders) (Figure 7 C&D), suggesting functionally active BK channels are required for YM-25890-induced attenuation of bladder contractility.

Collectively, these results suggest that GqPCR activation promotes DSM excitability in part through PKC-dependent suppression of BK channel activity.

## DISCUSSION

The role of the urinary bladder appears remarkably straightforward, to store and release urine, yet how the sensation of fullness is perceived and conveyed to the central nervous system is not fully understood. While local phasic contractions produce pressure fluctuations within the bladder that correspond with bursts of afferent nerve activity, the underlying mechanism of these rhythmic contractions remains unclear. This creates a major knowledge gap, where an incomplete understanding of urinary bladder physiology limits our ability to target novel pathways that may contribute to bladder dysfunction. Approximately 15-40% of the United States population suffers from urinary bladder pathologies and treatment options are limited (Reynolds *et al*., 2016). Gaining insight into DSM excitability and contractility is crucial to understanding bladder disease progression.

The aim of this study was to determine whether GqPCR, which influence ion channels and contractility in many types of smooth muscle, regulate urinary bladder phasic contractility. Here we demonstrate that GqPCR activity in whole bladder preparations regulates contractility via modulation of DSM V_M_. Loss of GqPCR activity hyperpolarizes DSM to decrease action potentials, thereby decreasing Ca^2+^ signaling prevalance and phasic contractility. Additionally, we show that downstream effects likely occur through a PKC dependent mechanism, most probably involving BK channels.

GPCR are well known regulators of smooth muscle tissue contractility. The urinary bladder relaxes during filling, at least in part through a GsPCR mechanism secondary to beta adrenergic receptor activation, while GqPCR activation induces voiding contractions following stimulation of muscarinic receptors (Andersson & Arner, 2004).

Although GqPCR activity increases IP3 and DAG to activate IP3 receptors and PKC, respectively, action potentials and contraction of DSM are predominantly reliant on the influx of extracellular Ca^2+^ (Hashitani & Brading, 2003b; Hashitani *et al*., 2004; Nausch *et al*., 2010). With PKC activity known to inhibit K^+^ channels and activate Ca^2+^ channels, the two major types of ion channels governing V_M_ and DSM action potentials, we decided to examine the role of GqPCR on phasic contractility. Due to the ubiquitous expression of GqPCR in the urinary bladder, we investigated the effects of broad inhibition on bladder function using the YM-254890 compound.

Broad inhibition of GqPCR by YM-254890 markedly decreased phasic contractility (Figure 1C). In addition, using mice expressing a genetically encoded Ca^2+^ indicator in smooth muscle, we found that YM-254890 reduced Ca^2+^ signaling prevalence and also impaired the synchronized spread of Ca^2+^ signals across the bladder wall (Figure 4, Movies 1 & 2). We noted some areas of the bladder continue to initiate Ca^2+^ signaling activity in the presence of YM-254890, though the ability to synchronize with adjacent DSM cells and produce phasic contractions was severely impaired (Movie 1). These actions of YM-254890 are not likely attributable to direct modulation of Ca^2+^ entry through VDCC as YM-254890 had no effect on KCl-induced bladder contractions (Figure 3D). This suggests that GqPCR activity in DSM may not function as a “pacemaker” of phasic contractility per se but rather acts as an amplifier of Ca^2+^ signals. This observation could be leveraged to enhance or depress afferent signaling in cases of overactive or underactive bladder.

The open probability of VDCC increases as smooth muscle V_M_ depolarizes (Nelson *et al*., 1990). A mechanism by which the GqPCR signaling cascade could control DSM phasic contractility is through effects on V_M_, altering the open probability of VDCC. Hyperpolarization of DSM V_M_ decreases action potential frequency (Heppner *et al*., 1997; Hashitani & Brading, 2003b). Our microelectrode experiments demonstrated a mean hyperpolarization of 6 mV in DSM cells in response to GqPCR inhibition with YM- 254890 (Figure 6E); thus our results indicate GqPCR activity is integral to DSM V_M_ and the generation of TPE. Further, it is well known that PKC can regulate K^+^ channels in many types of smooth muscle, including urinary bladder. Here, we observed DSM V_M_ depolarization following activation of PKC with phorbol ester in the presence of YM- 254890, a result supported by Hristov et al. who observed PKC inhibited BK channel currents and depolarized isolated DSM cells in guinea (Hristov *et al*., 2014).

In addition to BK channels, Kv and SK channels contribute to DSM hyperpolarization (Hashitani & Brading, 2003a, b). After eliminating the K^+^ electrochemical gradient by exposing the bladder to 120 mM extracellular KCl, which depolarizes DSM V_M_, we found no effect on contractility following inhibition of GqPCR activity with YM-254890 (Figure 5). This suggests the effect of GqPCR activity is likely through a K^+^ channel mediated hyperpolarization, a result supported by our electrophysiology results. PKC can affect Ca^2+^ channels (Fish *et al*., 1988; Navedo *et al*., 2005). It has also been suggested that unknown cation channels in the urinary bladder could provide the initial depolarization required to activate VDCC (Malysz & Petkov, 2020), a possibility we cannot exclude based on our results. Following BK channel inhibition, we show YM-254890 was unable to eliminate TPE (Figure 7C), suggesting BK channels play a role in GqPCR mediated phasic contractility, though Ca^2+^, Kv, and/or SK channels may well be involved.

An important question that this study does not address is: what is the source of GqPCR activation? Due to the enormity of potential ligands, we chose to focus on the broad role of GqPCR in DSM contractions. Further complicating efforts to identify ligands is the potential for multiple GqPCR pathways converging to govern contractility, leading to compensatory effects. Because neither atropine (Figure 2) or tetrodotoxin inhibit phasic contractions in urinary bladder preparations, the source of GqPCR activation appears to originate from some cellular source within the bladder (Drumm *et al*., 2026). Prostaglandin E_2_, prostaglandin F2-alpha, thromboxane, serotonin, endothelin, bradykinin, neurokinin, and histamine are coupled to GqPCR and have been found to induce DSM contraction and/or amplify phasic contractility, though some of these receptors may function primarily in pathological states (Palea *et al*., 1998; Kim *et al*., 2002; Kajioka *et al*., 2004; Lee *et al*., 2007; Grundy *et al*., 2018; Stromberga *et al*., 2020a, c; Borsodi *et al*., 2021). One potential class of ligands that warrant further investigation, prostanoids, have been implicated in overactive bladder pathology and can increase phasic contractions (Dobrek & Thor, 2015; Stromberga *et al*., 2020b). We previously demonstrated inhibition of prostanoid synthesis via indomethacin decreased TPE and afferent nerve activity, while Jones et al. showed indomethacin and a cocktail of prostanoid GqPCR antagonists can inhibit agonist induced increases in phasic contractions from the mast cell degranulator compound 48/80 (Jones *et al*., 2023; Heppner *et al*., 2024). Additionally, Parajuli et al. observed prostaglandin E_2_ inhibited spontaneous transient BK currents and enhanced DSM phasic contractility while the thromboxane receptor is a potent agonist of DSM contractility (Parajuli *et al*., 2014; Stromberga *et al*., 2020b). Another intriguing prospect is that GqPCR are activated by stretch during bladder filling (Wellner & Isenberg, 1993a, b). There are multiple reports of potential mechanosensitive GqPCR, and the forces generated during bladder filling are a plausible source for GqPCR activation (Shetty *et al*., 2025). Further, multiple compounds that could influence contractility, including ACh, ATP, and prostanoids, are released during bladder filling (Dobrek & Thor, 2015; Merrill *et al*., 2016). Either direct mechanical activation, stretch induced ligand production, or both appear to be prime candidates in modulating phasic contractility, and warrant further study.

Future efforts should focus on determining (1) which GqPCR(s) regulate DSM phasic contractility (2) how GqPCR(s) are activated (3) which ion channels are involved and (4) if there are additional GqPCR effectors (i.e. IP3 or Rho Kinase pathways) involved in modulating DSM contractility. These questions will help develop new models of bladder pathologies and provide additional drug targets for the treatment of overactive and underactive bladder.

## METHODS

### Animals

*Ethical Approval.* All experimental protocols were reviewed and approved by the Institutional Animal Care and Use Committee of the University of Vermont. Mice were group-housed on a 12-hour light/dark cycle with ad libitum delivery of food and water and were handled in accordance with ARRIVE guidelines. Urinary bladders were isolated from male and female C57Bl6/J mice (The Jackson Laboratory, stock number 000604; Bar Harbor, ME, USA). Mice were euthanized by intraperitoneal injection of a lethal dose of euthanasia solution, followed by decapitation. Urinary bladders were removed and placed in ice-cold dissection solution (see *Reagents and Solutions*). *Ex-Vivo Pressurized Bladder.* The ex-vivo pressurized bladder was performed as described previously (Heppner *et al*., 2024). In brief, after mice were euthanized, the urinary bladder was removed and placed in dissection solution. Both ureters were isolated and tied off with 5/0 silk suture near the bladder wall to prevent leaking of intravesical PSS and loss of pressure within the bladder. The preparation was moved to a custom fabricated recording chamber, cannulated via the urethra, and attached to a syringe pump (Model 4400-001, Harvard Apparatus) and an in-line pressure transducer (PT-F, Living Systems Instrumentation, Saint Albans, Vermont USA) that was connected to a signal conditioner set to 100 mV/cmH_2_O (Model NL-108, Digitimer, Hertfordshire, UK). This setup allows continuous infusion of PSS into the bladder and simultaneous recording of intravesical bladder pressure. The recording chamber was superfused with PSS maintained at 37°C and pH maintained at 7.4 by bubbling the solution with 20% O_2_/5% CO_2_/75% N_2_. To generate transient pressure events, the bladder was cannulated and multiple successive filling/emptying cycles were performed to obtain stable urodynamic profiles. Based on cystometry recordings of C57BL/6J mice which demonstrate that voiding threshold occurs around 11 mmHg and peak contraction pressures occur at 26 mmHg (Herrera2003), the bladder was then partially filled with PSS at a rate of 30 μL/min to induce TPE (6-12 mmHg) and allowed to stabilize over ∼20 minutes. During this time TPEs stabilize in frequency and amplitude. Data acquisition was performed using a Power3A analog-to-digital converter and Spike2 software (Cambridge Electronic Design, Cambridge UK) at a rate of 100 samples per second. TPE and Ca^2+^ imaging (8000 frames) were quantified over a 5-min isovolumetric period using peak detection features in Spike2 software (apply smoothing to pressure signal with a time constant of 0.5 sec followed by Spike2 “Peak Find” function with an amplitude threshold of 0.05 mmHg).

*Ca^2+^ Imaging Studies.* Bladders were isolated from B6-GC6f x SMMHC-CreER^T2^ or Cdh5-GCaMP8 x Acta2-RCaMP. The B6-GC6f x SMMHC-CreER^T2^ mice were generated from *Myh11*-CreER^T2^-RAD mice (JAX Stock no:037685) and crossed with mice harboring floxed GCaMP6f (Ai95D; Jax Stock no: 028865) to express green fluorescent protein in smooth muscle cells. The Cdh5-GCaMP8 x Acta2-RCaMP mice expressed red fluorescent protein in smooth muscle cells (acta2-RCaMP, stock #028345; Jackson Laboratory; CHROMus collaboration) (Ferris2023). No differences in fluorescence were noted between the two reporter mouse strains. The tissue bath was placed on the stage of a wide-field fluorescence microscope (Olympus MVX10) with a 1X PLANAPO objective (Olympus) to observe fluorescent Ca^2+^ activity. An LED excitation light source was used (XT720S, X-Cite Xylis) and a dichroic mirror filtered the excitation and emission light paths (ET470/40x, T495lpxr, ET525/50m, catalogue no. 49002). Fluorescence was captured with an Andor Zylya-4.2P CMOS camera (2048x2048x16bit) and micromanager 1.4 software (Edelstein 2010 & 2014) at 22.1 frames per second (fps). A data acquisition system (Spike2 software) collected the pressure signal and frame acquisition signal from the Zyla camera simultaneously at a rate of 100 samples per second. This approach resulted in a synchronized data stream for the acquisition of each video frame and intravesical pressure. Ca^2+^ imaging during analysis of TPE occurred over ∼5 minutes (8000 frames) for ex-vivo pressurized bladders.

#### Movie Preprocessing

The following fluorescent Ca^2+^ event analysis has been previously described (Herrera 2026) and described again in brief in the following sections. Ca^2+^ movies (8Gb) were imported into ImageJ and bladder cropped from non-bladder background (2-3Gb) and a debleaching and deflickering routine was performed to correct for any dimming or unstable light conditions. The recording was then corrected with an angular an XY dolly normalization to account for bladder movement during contractions.

#### Ca^2+^Extraction

Movies were then filtered (Gaussian Blur: 5 x 5 pixels, SD = 1.0) to reduce granular shot noise and no temporal filtering was used. A modified Standard Deviation of Quiescence (SDqe) routine (Herrera *et al*., 2026) was used to demarcate pixels in which fluorescence was elevated above background fluctuations in intensity (SD_min_ = 6.0, SD_threshold_ = 2.1 with a Quiescence Estimator (QE) of between 25%. Extracted particles were size filtered and saved as coordinate-based spatio temporal objects.

#### Ca^2+^ Event Refinement

Due to a greater volume at the outer edge of the bladder wall when projected on a plane and non-uniform Ca^2+^ events causing wall movement, ∼0.25mm of the perimeter were masked out leaving only extracted Ca^2+^ events from areas largely free from edge volume and wall motion artifacts. Ca^2+^ events evoking DSM contractions in large regions of the bladder can still produce spatial movement artifacts to occur. These distortion artifacts are small and could be filtered out without affecting DSM cell Ca^2+^ events. Any floating debris passing over the bladder during the recording was removed by deleting trajectories of particles observed beyond the edge of the bladder wall.

#### Ca^2+^ Event Measurements: Amplitude

We measured frequency and duration of Ca^2+^ events rather than amplitude because masking the outer edge of the bladder wall means we observe primarily DSM action potentials. Due to DSM action potentials having fairly uniform amplitudes, measurement of frequency, duration and area of Ca^2+^ events provides a more unbiased measurement of bladder contractility than amplitude per se.

#### Ca^2+^ Event Measurements: Prevalence

To best summarize the overall amount of Ca^2+^ activity throughout the entire visible hemisphere of isolated bladders during the 300 s recordings, a measure of Ca^2+^ event prevalence was calculated. This measure accumulates the area and duration of extracted Ca^2+^ events which can be mapped onto the bladder surface (Figure 4) or condensed into a single value by expressing prevalence in relation to the percent of bladder surface area (%Bladder area x s.min^-1^, See Movie 2).

#### Ca^2+^ Event Measurements: Coincidence

Prevalence measurements do not offer any insight into the synchrony or propagation of Ca^2+^ activity or if it is confined to individual DSM cells. We adapted Euclidian Distance Mapping (EDM) routines to radially scan within and around each pixel of each active Ca^2+^ event to gauge the proportion of nearby cells that were coincidently active. This provides a measure to determine if a Ca^2+^ event was isolated or spread across the bladder wall to produce a phasic contraction. By adapting Euclidian Distance Mapping (EDM) routines to radially scan 0.96mm within and around each pixel of each active Ca^2+^ event, we could gauge the proportion of nearby cells that were coincidently active at the exact same time as the reference activity. Slight delays in activation of neighboring DSM cells were accommodated due to most Ca^2+^ events having durations greater than 1 frame (44 ms). Raw coincidence results were normalized and are expressed as % of maximum search area activity within a radius of 1mm (%MSAA_r=1mm_). We choose to present coincidence as both the highest value at every point on the bladder surface (Max%MSAA_r=1mm_) and average coincidence (Avg%MSAA) to provide a visual reference for areas of the bladder expressing synchrony and the average amount of synchrony that occurred in these regions (Figure 4). Coincidence was mapped onto the bladder surface with fire coloring representing %MSAA ranging from 0 (no cells active at the same time) to 100 (all cells in the search radius active at the same time).

#### Membrane Potential Measurements

Urinary bladders were isolated and cut along the ventral wall from the bladder neck to dome and pinned flat in a dissection chamber. Smaller sheets were then dissected (∼0.5 mm x 1-2 mm) with intact urothelium and pinned detrusor smooth muscle side up in a recording chamber. Preparations were superfused with PSS warmed to 37°C with pH maintained at 7.4 by bubbling the solution with 20% O_2_/5% CO_2_/75% N_2_. Sharp microelectrodes were pulled from fire polished borosilicate microcapillary tubes (1.2 mm out diameter and 0.69 mm inner diameter, Sutter Instruments) to obtain tips with a resistance of ∼100-200 MΩ.

Microelectrodes were filled with 0.5 M KCl and iFlour-488 Hydrazide (3μM) and successful impalement of DSM cells was confirmed by z-stack imaging with an upright spinning disk confocal microscope (Andor). V_M_ recordings were made using an Axoclamp-2A digital amplifier and HS-2 headstage (Molecular Devices) with low-pass filtering (cutoff 1 kHz) and digitized and stored using Digidata 1322A and pCLAMP 9 software (Molecular Devices). Recordings used for analysis (Clampfit 11.4 software, Molecular Devices) demonstrated a steep negative deflection of V_M_ upon impalement, baseline V_M_ was stable for at least 5 min, and there was an immediate return to 0 mV once the microelectrode was removed.

#### Reagents and Solutions

Dissection solution consisted of 80 mM monosodium glutamate, 55 mM NaCl, 6 mM KCl, 10 mM glucose, 10 mM *N*-2- hydroxyethylpiperazine-*N’*-2-ethanesulfonic acid, 2 mM MgCl_2_, pH adjusted to 7.3 with NaOH. PSS contained 119 mM NaCl, 4.7 mM KCl, 24 mM NaHCO_3_, 1.2 mM KH_2_PO_4_, 2.5 mM CaCl_2_, 1.2 mM MgSO_4_, 11 mM glucose, and aerated with 20% O_2_/5% CO_2_/75% N_2_ to obtain pH 7.4. Euthanasia solution (Euthasol) was sourced from Midwest Veterinary Supply and contained pentobarbital sodium (390 mg/ml), phenytoin sodium (50 mg/ml), 10% ethanol, 18% propylene glycol, rhodamine B (0.003688 mg/ml), 2% benzyl alcohol (preservative) in water. The following pharmacological agents were used: YM-254890 and diltiazem hydrochloride (Tocris); phorbol 12,13-dibutyrate and paxilline (Cayman Chemical Company); and carbachol and atropine (Sigma-Aldrich).

## STATISTICS AND ANALYSIS

GraphPad Prism (GraphPad Software, Boston, MA) was used for all statistical analyses. Statistical significance was set at P < 0.05. T-tests, repeated measures ANOVA, or mixed-effect analysis were used to compare means as specified in figure legends. Summary data are presented as mean ± standard deviation (SD).

## DATA AVAILABILITY

All individual data used to generate figures, perform statistical analysis, and inform overall conclusions are found in respective summary data graphs. Detailed methodology to attain standard deviation based Ca^2+^ analysis and transient pressure analysis can be found in previously published articles (Herrera *et al*., 2026).

## AUTHOR CONTRIBUTIONS

JLR, GMH and NRK designed experiments and conceptualized the project. JLR, TJH, and SS conducted experiments. GWH developed Ca^2+^ analysis methodologies. JLR and GWH analyzed data. GMH, NRK, and MTN provided resources and animal models. JLR and GMH wrote the manuscript. TJH, GWH, NRK, and MTN provided feedback and edited the manuscript.

## ACKKNOWLEDGMENTS

The authors thank H. Ryan, H. Fallon, C. Dalton and N. Cashen for technical assistance and animal care.

## MOVIE LEGENDS

**Movie 1:** Example of Ca^2+^ events and spread along DSM bundles from raw fluorescence images. Left: Raw gray scale fluorescence at baseline, Right: Raw gray scale fluorescence from the same bladder following application of YM-254890.

**Movie 2:** Example of cumulative prevalence of Ca^2+^ events from the same bladder shown in Movie 1. Ca^2+^ prevalence value is depicted in both color spectrum and height for baseline conditions (Left) and after application of YM-254890 (Right).

